# Smoothness discriminates physical from motor imagery practice of arm reaching movements

**DOI:** 10.1101/2021.09.06.459053

**Authors:** Célia Ruffino, Dylan Rannaud Monany, Charalambos Papaxanthis, Pauline M. Hilt, Jérémie Gaveau, Florent Lebon

**Affiliations:** INSERM UMR1093-CAPS, Université Bourgogne Franche-Comté, UFR des Sciences du Sport, F-21000, Dijon

**Keywords:** motor imagery, movement smoothness, feedbacks, internal models, motor learning

## Abstract

Physical practice (PP) and motor imagery practice (MP) lead to the execution of fast and accurate arm movements. However, there is currently no information about the influence of MP on movement smoothness, nor about which performance parameters best discriminate these practices. In the current study, we assessed motor performances with an arm pointing task with constrained precision before and after PP (n= 15), MP (n= 15), or no practice (n= 15). We analyzed gains between Pre- and Post-Test for five performance parameters: movement duration, mean and maximal velocities, total displacements, and the number of velocity peaks characterizing movement smoothness. The results showed an improvement of performance after PP and MP for all parameters, except for total displacements. The gains for movement duration, and mean and maximal velocities were statistically higher after PP and MP than after no practice, and comparable between practices. However, motor gains for the number of velocity peaks were higher after PP than MP, suggesting that movements were smoother after PP than after MP. A discriminant analysis also identified the number of velocity peaks as the most relevant parameter that differentiated PP from MP. The current results provide evidence that PP and MP specifically modulate movement smoothness during arm reaching tasks. This difference may rely on online corrections through sensory feedback integration, available during PP but not during MP.

## Introduction

Motor skill learning is a central process in everyday life, sustaining adaptation and efficiency of motor behaviors in constantly changing environments. Through physical practice (PP), movements are performed faster, more accurately, and require less energy consumption (Willingham, 1998). Even if continuous and extended practice is known to greatly and durably improve motor performance (Robertson et al., 2004; Kitago and Krakauer, 2013), positive effects of practice can also be observed within a single training session. A growing number of studies indeed observed that a few minutes of practice is sufficient to induce gains in motor performance (e.g., speed and accuracy) on a wide variety of motor tasks, such as sequential finger-tapping or arm-reaching tasks (Karni et al., 1998; Walker et al., 2003; Gentili et al., 2006, 2010; Spampinato and Celnik, 2017; Ruffino et al., 2021). This fast learning process, known as motor acquisition, is considered as the first step towards the formation of new and robust motor memories.

Although skill learning usually requires PP, alternative forms of practice also exist. Among these, motor imagery, that is the mental simulation of an action without associated motor output, has been largely documented. In fact, mental practice (MP) improves several aspects of motor performance, such as movement accuracy, speed, and force (Yue and Cole, 1992; Yágüez et al., 1998; Ranganathan et al., 2004; Gentili et al., 2006, 2010; Allami et al., 2008; Lebon et al., 2010; Grosprêtre et al., 2018; Ruffino et al., 2021). Performance increases following MP is associated with specific neural mechanisms at both cortical and spinal levels (Avanzino et al., 2015; Grosprêtre et al., 2019; Ruffino et al., 2019). Specifically, an acute session of MP induces use-dependent plasticity into the primary motor cortex (Ruffino et al. 2019) and spinal circuitry (Grosprêtre et al. 2019). At the functional level, it is proposed that motor performance improvement following MP may reflect an update of internal forward models (Kilteni et al., 2018; Dahm and Rieger, 2019; Ruffino et al., 2021). Basically, internal forward models are neural network that predict the future sensorimotor state (e.g., velocity, movement duration, position) given the current state, the efferent copy of the motor command, and the goal of the movement (Kawato et al., 2003; Wolpert & Flanagan, 2001). Kilteni et al. (2018) strongly supported this assumption by showing that the sensory consequences of imagined movements are predicted during motor imagery.

Although PP and MP share common mechanisms, a number of dissimilarities also exist. Perhaps the main difference, at least the most visible, is that during MP there is no sensory feedback about movement (position velocity and acceleration), since the imagined segment in inert. In error-based motor learning process, external sensory feedback is necessary to update the prediction of the internal forward model, via the discrepancy between the predicted state and the actual state (Kawato et al., 2003; Shadmehr et al., 2010; Shadmehr & Krakauer, 2008; Wolpert et al., 2011). Better state prediction will, in turn, improves the controller and thus the motor output. In the case of a model-free motor learning process, external feedback directly improves the controller through reward predictions error (Criscimagna-Hemminger et al., 2010; Izawa and Shadmehr, 2011). The absence of external feedback during MP could explain why after PP the performance improvement is better than after MP (Ingram et al., 2019), but does not explain the underline mechanism. Theoretically, it is proposed that the difference between the prediction and the desired outcome based on stored movement representations would be returned as an input to improve the subsequent motor command via a “self-supervised process”, explaining motor performance improvement despite the external feedback during MP (Gentili et al., 2010, Ruffino et al. 2021).

Up to now, performance improvement after MP has been measured, and compared to PP, mainly on three parameters: force, accuracy, and speed. Nonetheless, other parameters are of importance for motor performance, such as movement smoothness that is enhanced after PP (Balasubramanian et al., 2015). Smoothness is related to the form of the velocity profile, which is singled-peaked with one acceleration and one deceleration phase. When the motor command is inaccurate a number of sub-movements are necessary to correct it, creating thus a non-optimal and clumsy movement (Kelso et al., 1979; Ketcham et al., 2002; Ketcham & Stelmach, 2004). Intriguingly, the effects of MP on this parameter are yet unknown.

In the current study, we sought to compare PP and MP, considering spatial, temporal, and smoothness parameters. We recorded movement-related trajectories on a graphic tablet from two training groups (PP and MP) and one control group (Ctrl, absence of practice). In line with the literature, we first hypothesized that PP and MP would similarly enhance arm reaching movements, with improvements for all parameters but with greater gains for PP. Alternatively, temporal parameters would similarly improve following PP and MP, as sensory feedback is not a prerequisite in that case, whereas spatial and smoothness parameters would be less improved after MP.

## Method

### Participants

Forty-five right-handed adults participated in this study after giving their informed consent. All were healthy and self-reported being free from neurological or physical disorders. The participants were randomly assigned into three groups: the Physical Practice group (PP, n = 15, 6 females, mean age: 25± 2 years old), the Mental Practice group (MP, n = 15, 9 females, mean age: 25 ± 6 years old), and the Control group (Ctrl, n = 15, 7 females, mean age: 28 ± 4 years old).

Motor imagery vividness of the MP group was assessed by the French version of the revised Movement Imagery Questionnaire ‘MIQr’ (Lorant and Nicolas, 2004). The MIQr is an 8-item self-report questionnaire, in which the participants rate the vividness of their mental images using 7-point scales ranging from 1 (really difficult to feel/visualize) to 7 (really easy to feel/visualize), on two modalities (visual imagery and kinesthetic imagery). The average score obtained in the current study was 42.1 ±9.5 (maximum score: 56; minimum score: 8), revealing good imagery capabilities.

### Experimental Device

The participants were comfortably seated on a chair in front of a graphic tablet (Intuos4, XL, Wacom, Krefeld, Germany), on which four square-targets (1×1 cm) were presented (see Fig.1A). The distance between the participants’ sternum and the first target (T1) was 10 cm. One trial included 10 successive target-to-target movements in the following order: 1 – 2 – 3 – 4 – 1 – 2 – 3 – 4 – 1 – 2 – 3. The participants were instructed to reach each target with a pencil as accurately and as fast as possible (Fig.1B).

**Fig. 1.**
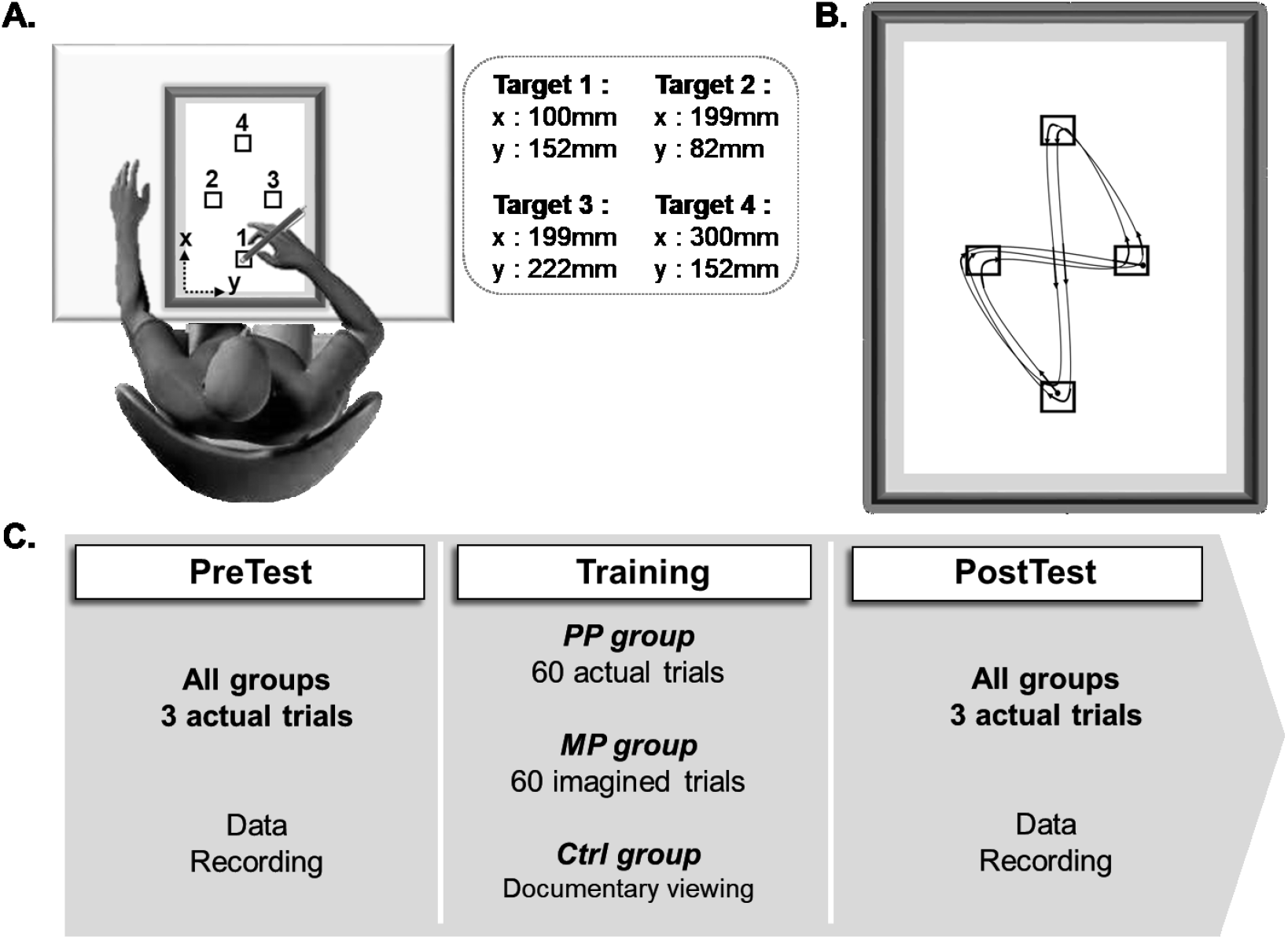
***A.*** Participants’ position and targets location on the graphic tablet. ***B.*** One trial included 10 successive track movements between the targets, as fast as possible without missing any target (1 – 2 – 3 – 4 – 1 – 2 – 3 – 4 – 1 – 2 – 3). ***C.*** Experimental procedure. The protocol included 2 test sessions of 3 actual trials (data recording for each trial) and one training session. During the training session, the Physical Practice (PP) group physically executed 60 repetitions of the 10 movements, and the Mental Practice (MP) group mentally simulated 60 repetitions of the 10 movements. The Control (Ctrl) group watched a non-emotional documentary.

### Experimental procedure

The experimental protocol included two *test* sessions (PreTest and PostTest) and one *training* session (Fig. 1C). During the *test* sessions, all the participants performed 3 actual trials as fast and accurately as possible. During the training session, the participants of the PP were trained as fast and accurately as possible to the task, while those of the MP group were instructed to imagine themselves performing the task as fast and accurately as possible, combining kinesthetic and visual (first-person perspective) imagery modalities. Both training groups performed 60 trials, divided into 6 blocks with 1-min rest between blocks to avoid mental fatigue (Rozand et al., 2016). The Ctrl group watched a non-emotional documentary (“Home”, directed by Y. Arthus-Bertrand, 2009) for 30 min (the approximate time of both training sessions).

### Kinematics recording and analysis

We recorded movement kinematics at 100Hz using a graphic tablet (Intuos4 XL, Wacom, Krefeld, Germany). The spatial resolution in the present experiment was less than 1 mm. Data processing was performed using custom programs written in Matlab (Mathworks, Natick, MA). Position signals in the horizontal plane (X, Y) were low-pass filtered using a digital fifth-order Butterworth filter (zero phase distortion, Matlab ‘butter’ and ‘filtfilt’ functions) at a cut-off frequency of 10 Hz.

We computed five parameters for each trial: i) movement duration (MD), i.e., the total time elapsed between the moment when the pencil exited the first target and entered the final target; ii) distance, i.e., the total two-dimensional displacement; iii) mean velocity (Vmean), i.e., the average inter-target movement speed; iv) maximal velocity (Vmax), i.e., the average of maximal inter-target movement speed; and v) number of velocity peaks (NbPeaks), i.e., the number of local maxima detected on velocity profiles. We used this parameter to quantify movement smoothness (Brooks et al., 1973; Fetters and Todd, 1987; Balasubramanian et al., 2015); the smaller the number of peaks, the smoother the movement.

For each parameter, we calculated the gain between PreTest and PostTest. To systematically represent gains with positive values, we calculated gains for MD, NbPeaks, and distance as follows:

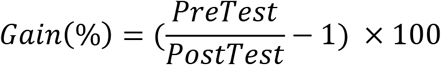

and for Vmax and Vmean as follows:

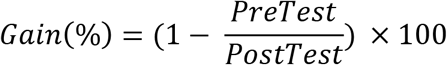

### Electromyographic recording and analysis

To verify that muscles were not activated during mental training (MP group), electromyographic (EMG) activity of the biceps brachii (BB) and the triceps brachii (TB) muscles of the right arm were recorded during each imagined trial and compared to EMG activity at rest (10-second recording before training). We used pairs of bipolar silver chloride circular (10-mm diameter) surface electrodes. We positioned the electrodes parallel to muscle fibers, over the middle of the muscles belly with an inter-electrode (center-to-center) distance of 20 mm. The reference electrode was positioned on the medial elbow epicondyle. After shaving and dry-cleaning the skin with alcohol, the impedance was below 5 kΩ. EMG signals were amplified (gain 1000), filtered (with a bandwidth frequency ranging from 10 Hz to 1 kHz), and converted for digital recording and storage with PowerLab 26T and LabChart 7 (AD Instruments). We analyzed the EMG patterns of the muscles by computing their activation level (RMS, root mean square) using the following formula:

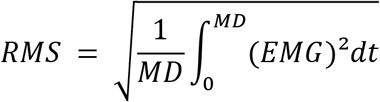

### Statistical analysis

We performed the analyses on motor gains to reduce variability between participants, especially at PreTest. We primarily checked the normality of the data (Shapiro-Wilk test), the equality of variance (Levene’s test), and the sphericity (Mauchly’s test).

First, we used unilateral one-sample t-tests compared to the reference value 100 to check whether motor performances improved between Pre and PostTest, for each parameter and each group. Cohen’s d was reported for each test and the statistical significance threshold was set at 0.05. P values were adjusted using the Bonferroni method (divided by the number of comparisons).

To compare gains between groups, we then performed one-factor ANOVAs with *Group* as a between-subject factor and planned comparisons using orthogonal contrasts analysis for each parameter (Howell, 2012). We constructed a contrast matrix to test the following *a priori* assumptions: i) MP and PP led to better gain when compared to an absence of practice, i.e., Ctrl group (contrast C1), and ii) PP led to better gain when compared to MP (contrast C2).

To identify the parameters that best discriminated the groups, we finally realized a stepwise generalized linear discriminant analysis. This exploratory data analysis first consisted in the identification of discriminant parameters, and then in the creation of functions that combined these discriminant parameters. The resulting functions were used as a linear classifier to investigate the data organization according to the categorical predictors (i.e., groups) and the independent variables (i.e., parameters). To test if the discriminant functions classified the experimental observations in their respective groups better than chance (i.e., if the combinations of identified factors were indeed relevant to group discrimination), we used the Press Q statistic (Hair et al., 1998).

Also, to ensure that participants of the MP group did not activate their muscles during MP, we used Friedman’s ANOVAs, comparing the EMG activity of each imagined block with the rest condition, for each muscle (BB and TB).

## Results

### Summary data

Table 1 reports the mean values and the mean gains for the five kinematic parameters.

**Table 1.** Average values and standard deviation (SD) for the PreTest, PostTest, and gains of the five parameters and the three experimental groups. MD: Movement duration; Vmean: Mean velocity; Vmax: Maximal velocity; NbPeaks: Number of peaks, s: second, cm: centimeter.

|  |  | PreTest | PostTest | Gain (%) |
| --- | --- | --- | --- | --- |
|  |  | Mean (SD) | Mean (SD) | Mean (SD) |
| PP | MD (s) | 4.87 (0.97) | 4.17 (0.78) | 17.31 (15.26) |
|  | Distance (cm) | 162.01 (5.35) | 163.19 (6.61) | -0.61 (4.23) |
|  | Vmean (cm/s) | 32.61 (8.07) | 37.79 (8.42) | 13.31 (11.81) |
|  | Vmax (cm/s) | 71.98 (22.12) | 81.37 (26.18) | 10.39 (10.34) |
|  | NbPeaks | 5.63 (2.14) | 4.18 (1.47) | 37.07 (29.37) |
| MP | MD (s) | 4.80 (0.92) | 4.43 (0.82) | 9.11 (13.16) |
|  | Distance (cm) | 164.42 (5.24) | 164.23 (3.72) | 0.1 (1.94) |
|  | Vmean (cm/s) | 32.43 (5.38) | 35.25 (5.91) | 7.38 (11.34) |
|  | Vmax (cm/s) | 73.44 (13.25) | 81.25 (12.35) | 9.39 (11.43) |
|  | NbPeaks | 4.60 (1.13) | 3.87 (0.77) | 18.62 (15.25) |
| Ctrl | MD (s) | 4.87 (0.84) | 4.67 (0.87) | 4.85 (7.89) |
|  | Distance (cm) | 163.55 (4.34) | 163.54 (4.44) | 0.02 (1.49) |
|  | Vmean (cm/s) | 32.31 (5.94) | 33.76 (6.74) | 3.82 (6.53) |
|  | Vmax (cm/s) | 71.59 (14.27) | 71.18 (15.56) | -1.03 (6.30) |
|  | NbPeaks | 6.22 (2.27) | 5.75 (2.45) | 11.48 (19.15) |

### One sample t-tests

Firstly, to check whether motor performances improved between PreTest and PostTest after practices, we used unilateral one-sample t-tests compared to the reference value 100. Table 2 reports the results and effect sizes for one-sample t-tests analysis.

**Table 2.**
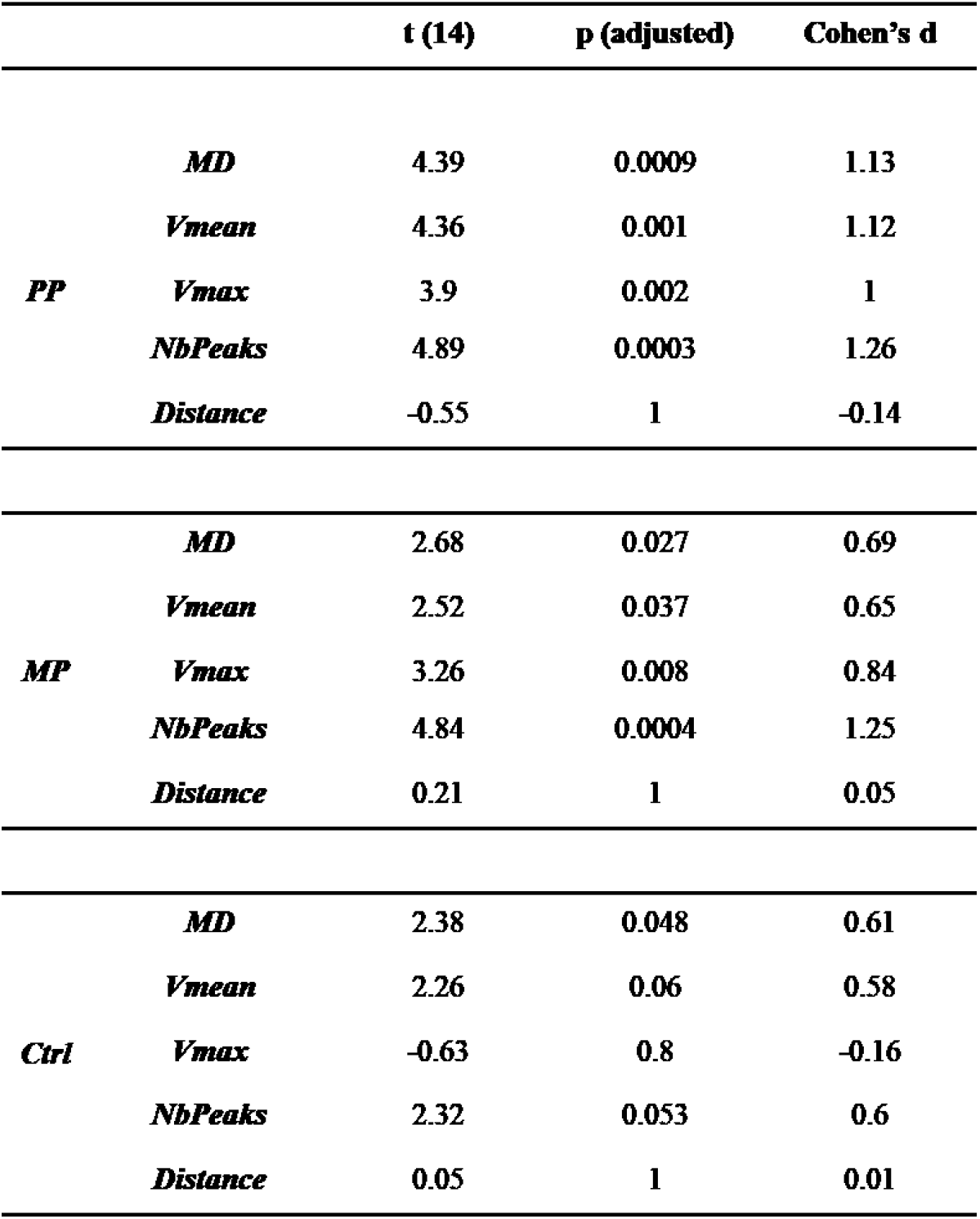
Summarized results and effect sizes for one-sample t-tests analysis. PP: Physical practice group; MP: Mental practice group; Ctrl; Control group. MD: Movement duration; Vmean: Mean velocity; Vmax: Maximal velocity; NbPeaks: Number of peaks.

PP and MP groups significantly improved their performances between PreTest and PostTest sessions. Precisely, MD and NbPeaks decreased, while Vmax and Vmean increased after practice. The Ctrl group improved movement duration. No significant effect of Distance was observed (all *p’s* > 0.05).

### One factor ANOVA and contrast analysis

Secondly, we compared gains between groups by means of one-factor ANOVAs and planned comparisons. The results are depicted in Figure 2.

**Fig. 2.**
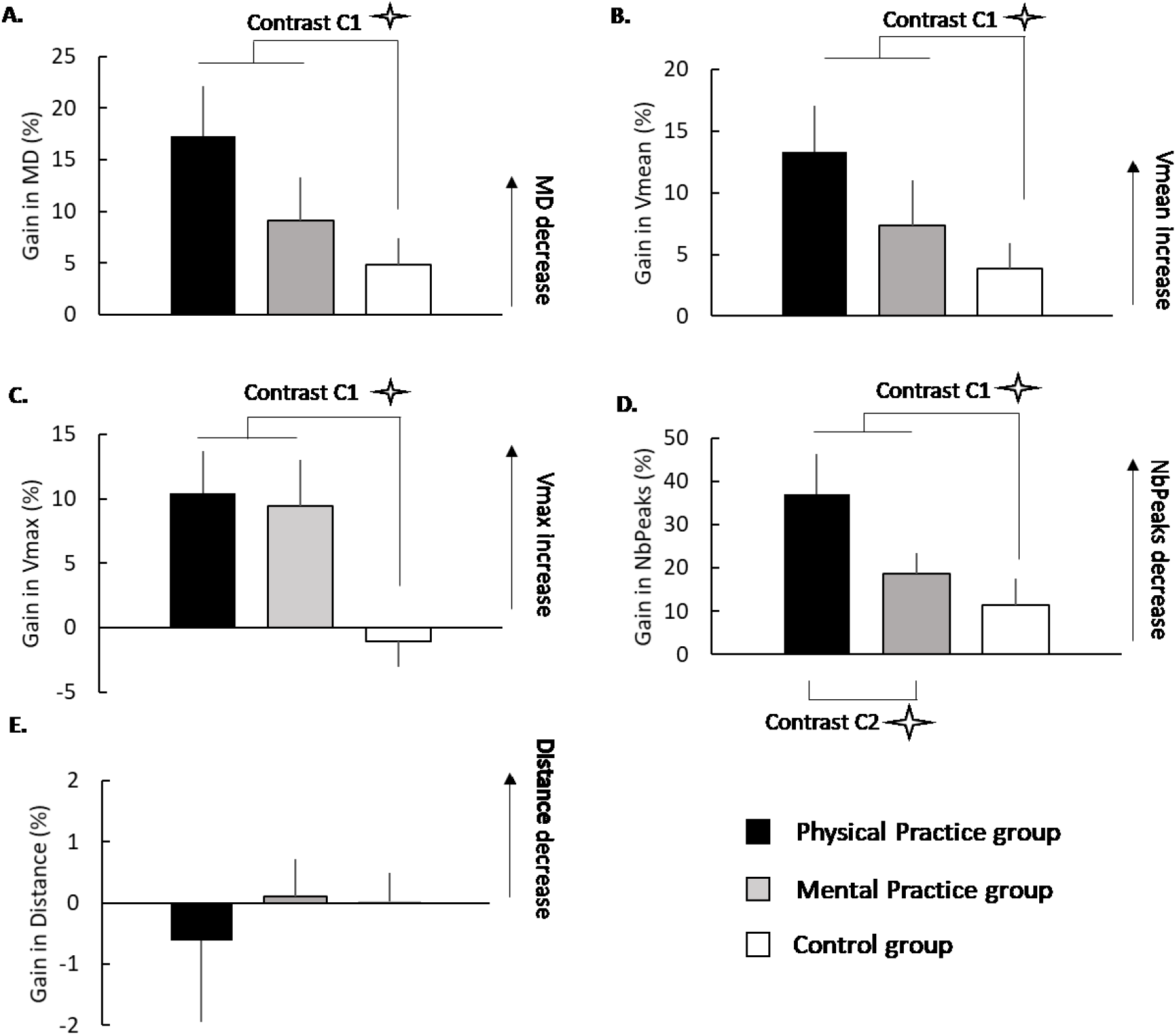
Mean gains (+SE) for MD (***A.*** movement duration), Vmean (***B***. mean velocity), Vmax (***C.*** maximal velocity), NbPeaks (***D.*** number of peaks) and in distance (***E***.) for each group. Stars indicate significant effect of the contrasts (C1: Ctrl vs PP + MP and C2: PP vs MP).

The ANOVA revealed a main effect of Group for MD (*F*_2, 42_ = 3.85, *p*= 0.03, *η_p_^2^*= 0.15), Vmean (*F*_2, 42_ = 3.32, *p*= 0.045, *η_p_^2^*= 0.14), Vmax (*F*_2, 42_ = 6.64, *p*< 0.01, *η_p_^2^*= 0.24) and NbPeaks (*F*_2, 42_ = 5.37, *p*< 0.01, *η_p_^2^*= 0.2), but nor for Distance (*F*_2, 42_ = 0.29, *p*= 0.75). The contrast analysis revealed significant effect of the contrast C1 (Ctrl vs MP + PP) for MD (*t*= 2.11, *p*= 0.04, *Cohen’s d*= 0.69), Vmean (*t*= 2.02, *p*= 0.049, *Cohen’s d*= 0.66), Vmax (*t*= 3.63, *p*< 0.01, *Cohen’s d*= 1.18) and NbPeaks (*t*= 2.36, *p*= 0.02, *Cohen’s d*= 0.78), without effect for Distance (t= −0.3, *p*= 0.76). This confirms that, except for Distance, practice (MP & PP) improved motor performance when compared to the absence of practice (Ctrl). The contrast C2 (MP vs PP) revealed no statistically significant difference for MD, Vmean, Vmax or Distance (*all p’s* > 0.12). However, there was a significant effect for the contrast C2 regarding NbPeaks (*t=* 2.27, *p*= 0.03, *Cohen’s d*= 0.82), suggesting better performance after PP than MP.

### Stepwise generalized linear discriminant analysis

Finally, we performed a stepwise generalized linear discriminant analysis to identify the parameters that best discriminated the groups. The three groups (PP, MP, and Ctrl) were considered as the dependent variable and the gains for each parameter as the independent variable. The discriminant power of each variable was tested using a forward stepwise approach, revealing that Vmax and NbPeaks significantly contributed to group discrimination (*F*_2_ = 5.19, *p*< 0.01 and *F*_2_ = 3.99, *p*= 0.025, respectively), whereas MD (*F*_2_ = 1.41, *p*= 0.25), distance (*F*_2_ = 0.04, *p*= 0.96), and Vmean (*F*_2_ = 1.38, *p*= 0.26) did not.

The discriminant analysis gave two canonical functions (*Wilks’ λ*= 0.64; *χ^2^* (4) = 18.62, *p*< .01 for the first function; *Wilks’λ* = 0.84; *χ^2^* (1) = 7.43,*p*< .01 for the second one). The first discriminant function accounted for 30.94% of total variance, while the second one for 19.61%, for a total of 50.55%. The classification accuracy of the discriminant functions was 62.2% with a significant Press Q score (*χ^2^* (1) = 16.9, *p*< .01). This result ensures that the discriminant functions classified experimental observations in their respective groups better than chance (i.e., 50%). To conclude, the results of discriminant analysis and contrasts analysis suggest that Vmax discriminates practice from the absence of practice, whereas Nbpeaks also discriminates the performance improvement between PP and MP.

### Electromyographic analysis

Participants did not activate their muscles during mental training in comparison to rest. Statistical comparison (Friedman’s Anova) of EMG activity between each block of MP and rest revealed no significant difference neither for the BB muscle (*X^2^* =5.47; *p*= 0.48) nor for the TB muscle (*X^2^* = 10.39; *p*= 0.11).

## Discussion

In the current study, we identified the number of velocity peaks, an indicator of movement smoothness, as the most relevant parameter that differentiated PP from MP for an arm pointing task. While classical parameters as movement duration or maximal and mean velocity improved in a comparable extent following both practices, movement smoothness improved following MP but to a lower extent than that after PP. These findings provide relevant information about the specific influence of practice types on motor performance parameters.

### General motor performance improvement

Motor performance improvement of arm reaching movement have been widely investigated, considering both PP and MP (Gentili et al. 2006; Gentili et al. 2010; Yàgüez et al. 1998; Ingram et al. 2019). Our findings corroborate previous results, showing an improvement of temporal parameters, such as movement duration, mean and maximal velocity after both practices, compared to the Control group.

The improvement of motor performance following MP could be explained by the concept of forward internal model (Gentili et al. 2006; Gentili et al. 2010; Dahm et Rieger 2019; Kilteni et al. 2018). The internal model theory postulates the existence of predictive processes that simulate sensory prediction and the dynamic consequences of action (Wolpert and Flanagan, 2001; Friston, 2011; Kilteni et al., 2018). During the mental simulation of a movement, an efferent copy generated by the controller would be transmitted to predictive models, allowing to generate two predictions: a prediction of the consequences of action (Ruffino et al. 2021) and a prediction of the related-movement sensory afferents (Grush, 2004). The comparison between these predictions and the stored movement representation would be fed back as input to the controller, leading to an improvement of motor command despite the absence of feedbacks. Interestingly, PP and MP increased mean and maximal velocity to the same extent, but PP greater decreased the number of velocity peaks. These findings provide evidence that PP and MP may improve specifically the parameters of performance for arm reaching tasks.

### Movement smoothness discriminates physical and mental practices

The arm reaching movements can be decomposed in two distinct phases: i) an initial impulse phase, involving predictive loops and ii) a final phase, known as the corrective phase, implying online movement corrections (Elliott et al., 2001; Thompson et al., 2007). Kinematic analyses revealed that the first phase can be characterized by one or two high velocity peaks, permitting to quickly get closer to the target, while the second contains low secondary velocity peaks, which are likely to represent corrective sub-movements when approaching the target (Novak et al., 2002). The authors also suggested that PP leads to faster and precise initial movements in order to quickly approach the target and to reduce the number of corrective sub-movements, respectively. Here, we discuss the difference in number of velocity peaks between PP and MP groups, considering the insights of studies that investigated the absence of feedbacks during actual reaching movements (Khan et al., 2003; Franklin et al., 2017). These studies showed that the actual execution of fast and smooth movements during the initial phase is possible with or without feedbacks, whereas the reduction of the endpoint variability during the final phase is feedback-dependent. Because of the similar increase of Vmax for both groups and the decrease of NbPeaks, even for MP, we suggest that both PP and MP may lead to the execution of faster and precise initial movements that minimize the corrective phase and thus reduce the number of sub-movements. The distinction between PP and MP could thus stand in the corrective phase, where sensory feedbacks are necessary. Indeed, the feedbacks of actual movements during PP may help to optimize the corrective phase when approaching the target, and therefore to greater reduce NbPeaks in comparison to MP. The absence of sensory feedbacks during imagined movements would be an obstacle to reduce the number of sub-movements when actually approaching the target.

## Conclusion

In conclusion, the present study provided the first evidence that MP increased smoothness of arm-reaching movement, and that this performance parameter discriminated between PP from MP. Although, no sensory feedbacks are present during imagined movements, the increase of movement velocity would lead to greater smoothness after MP. Further studies could analyse a broader range of movements and tasks (e.g., to perform and/or imagine the movement at different velocities) to better understand the influence of MP on movements parameters.

## Authors contributions

CR, CP, and FL designed the experiment; CR and DRM recorded the data; DRM and JG analysed the data; CR and DRM developed figures; CR, DRM, CP and FL wrote the manuscript; CP, JG, PMH and FL provided feedback on the manuscript; all co-authors read and approved the submitted version.

